# Structure-guided disruption of the pseudopilus tip complex inhibits Type II secretion in *Pseudomonas aeruginosa*

**DOI:** 10.1101/303123

**Authors:** Jerry Yichen Zhang, Frédérick Faucher, Wenwen Zhang, Shu Wang, Nolan Neville, Keith Poole, Jun Zheng, Zongchao Jia

## Abstract

*Pseudomonas aeruginosa* utilizes the Type II secretion system (T2SS) to translocate a wide range of large, structured protein virulence factors through the periplasm to the extracellular environment for infection. In the T2SS, five pseudopilins assemble into the pseudopilus that acts as a piston to extrude exoproteins out of cells. Through structure determination of the pseudopilin complexes of XcpVWX and XcpVW and function analysis, we have confirmed that two minor pseudopilins, XcpV and XcpW, constitute a core complex indispensable to the pseudopilus tip. The absence of either XcpV or -W resulted in the non-functional T2SS. Our small-angle X-ray scattering experiment for the first time revealed the architecture of the entire pseudopilus tip and established the working model. Based on the interaction interface of complexes, we have developed inhibitory peptides. The structure-based peptides not only disrupted of the XcpVW core complex and the entire pseudopilus tip *in vitro* but also inhibited the T2SS *in vivo*. More importantly, these peptides effectively reduced the virulence of *P. aeruginosa* towards *Caenorhabditis elegans*.

---

*Pseudomonas aeruginosa* is a Gram-negative bacterium known for its severe pathogenicity. Therapeutic treatment of *P. aeruginosa* is difficult due to its low outer member permeability, enzymatic modification and efflux of antibiotics(Breidenstein, de la Fuente-Nunez et al., 2011). Its ability to catabolize a wide range of organic molecules guarantees its high adaptability and prevalence in ubiquitous environments(Boles & Singh, 2008, Filipiak, Sponring et al., 2012), which makes it one of the most predominant nosocomial pathogens. As an opportunistic pathogen, it tends to preferentially colonize in immunocompromised patients like those suffering from cystic fibrosis, cancer, or AIDS(Lore, Iraqi et al., 2015), and cause pneumonia, urinary tract infection and gastrointestinal infection(Beveridge & Kadurugamuwa, 1996, Hardalo & Edberg, 1997, Sawa, 2014, Young & Armstrong, 1972). As a result, the World Health Organization has recently deemed *P. aeruginosa* a Priority 1 pathogen that is in urgent need of new antibiotics(2017).

To promote infection, *P. aeruginosa*, like other pathogenic Gram-negative bacteria, has evolved complex secretion systems to mediate the passage of various macromolecular virulence factors across the cellular membrane. The Type II secretion system (T2SS) adopts a two-step translocation to transport large, structured exoproteins through the periplasm into extracellular milieu. The secreted virulence factors degrade and invade host tissues and cells, and facilitate bacterial survival and proliferation(Korotkov, Sandkvist et al., 2012, Ribet & Cossart, 2015). In *P. aeruginosa*, this sophisticated system, consisting of twelve Xcp proteins, is subdivided into four subassemblies: 1) the inner membrane platform composed of XcpP, XcpS, XcpY and XcpZ, 2) XcpR-hexamerized ATPase, 3) the outer membrane channel formed by XcpQ, and 4) the pseudopilus, featuring a piston-like structure (**Fig EVS1**)(Douzi, Ball et al., 2011). The pseudopilus, a homolog of the Type IV pili(Ayers, Howell et al., 2010, Peabody, Chung et al., 2003), serves as the central structure of the T2SS(Douzi, Filloux et al., 2012). It consists of five pseudopilins, in which four minor pseudopilins (XcpU, -V, -W and -X) form a quaternary pseudopilus tip complex, which is critical for the recognition and binding of secretion substrates(Douzi et al., 2011), atop the pseudopilin body polymerized by the major pseudopilin, XcpT.

Pioneered by Hol’s group, characterizations of pseudopilin-related crystal structures have provided incisive insights into the organization and function of the pseudopilus(Korotkov & Hol, 2008, Korotkov, Johnson et al., 2011, Yanez, Korotkov et al., 2008a, Yanez, Korotkov et al., 2008b), and the assembly, dynamics and secretion mechanisms have been gradually revealed(Cisneros, Bond et al., 2012a, Cisneros, Pehau-Arnaudet et al., 2012b, Nivaskumar, Santos-Moreno et al., 2016). It has been established that XcpV^(GspI)^and XcpW^(GspJ)^ significantly influence toxin secretion(Cisneros et al., 2012a, Franz, Douzi et al., 2011, Nivaskumar et al., 2016). XcpV acts as a nucleator that recruits other minor pseudopilins for pseudopilus tip assembly(Douzi, Durand et al., 2009). The loss of XcpV^(GspI)^ leads to the failure in the formation of the pseudopilus(Cisneros et al., 2012a), which ultimately results in the non-functional T2SS. XcpW^(GspJ)^ interacts with XcpV^(GspI)^ to form a stable binary complex(Yanez et al., 2008a), which is also critical for exoprotein secretion(Franz et al., 2011). XcpX^(GspK)^ is considered to control the length of the pseudopilus(Durand, Michel et al., 2005), the deletion of which may influence the PulA secretion in *Klebsiella Oxytoca(Cisneros et al., 2012a, Nivaskumar et al., 2016)*. XcpU^(GspH)^ forms the bottom piece of the pseudopilus tip. Although it is reported to interact with the major pseudopilin XcpT^(GspG)^ via their N-terminal hydrophobic α-helices(Douzi et al., 2009), its role in the T2SS remains elusive because its deletion does not affect toxin secretions(Cisneros et al., 2012b). Virulence factors are first recruited by XcpP^(GspC)^ to the secretion pore(Douzi et al., 2011). Once secretion is initiated, XcpT^(GspG)^ polymerizes into the pseudopilus body that extends into the channel(Alphonse, Durand et al., 2010, Durand, Alphonse et al., 2011), pushing the virulence factors out of the cell.(Reichow, Korotkov et al., 2010, Yan, Yin et al., 2017). Therefore, the pseudopilus tip complex is an ideal target for potentially inhibiting the T2SS.

Based on aforementioned studies and our secretion assays using a well-established *Pseudomonas aeruginosa* transposon mutant library that has been widely accepted as an efficient tool of studying phenotypes in *P.aeruginosa*(Brown & Wright, 2016, Hoffman, D’Argenio et al., 2005, Jacobs, Alwood et al., 2003), we have confirmed that XcpV and -W form a core pseudopilin complex in the pseudopilus tip, which is indispensable to T2SS functions. Through protein crystallography, we have obtained the structures of the XcpVW binary and XcpVWX ternary pseudopilin complexes. Combined with small-angle X-ray scattering (SAXS), we have for the first time revealed the architecture of the entire pseudopilus tip complex. Two inhibitory peptides were developed based on the interaction interface of XcpV and -W. These peptides destabilized and even disrupted the entire pseudopilus tip complex *in vitro*, which further inhibited the Type II secretion of typical virulence factors in *P. aeruginosa in vivo*. Moreover, these structure-based peptides were effective in attenuating *P. aeruginosa* infection in *Caenorhabditis elegans*, reducing the bacterial virulence, and increasing the survivability of the worms.

## Results

### Crystal structures of XcpVWX and XcpVW complexes

We determined the structures of the soluble forms of the minor pseudopilin complexes of XcpVWX and XcpVW (**Fig 1A, 1B and Table 1**) of *P. aeruginosa*. Instead of the previously reported refolding method(Korotkov & Hol, 2008), we used a thioredoxin tag to obtain soluble XcpX. The crystal structures of both complexes were determined at ∼2.0-Å resolution by molecular replacement. Although the overall folding is similar to homologous structures(Yanez et al., 2008a)’(Korotkov & Hol, 2008), our complex structures still feature some distinct structural characteristics.

**Figure 1.**
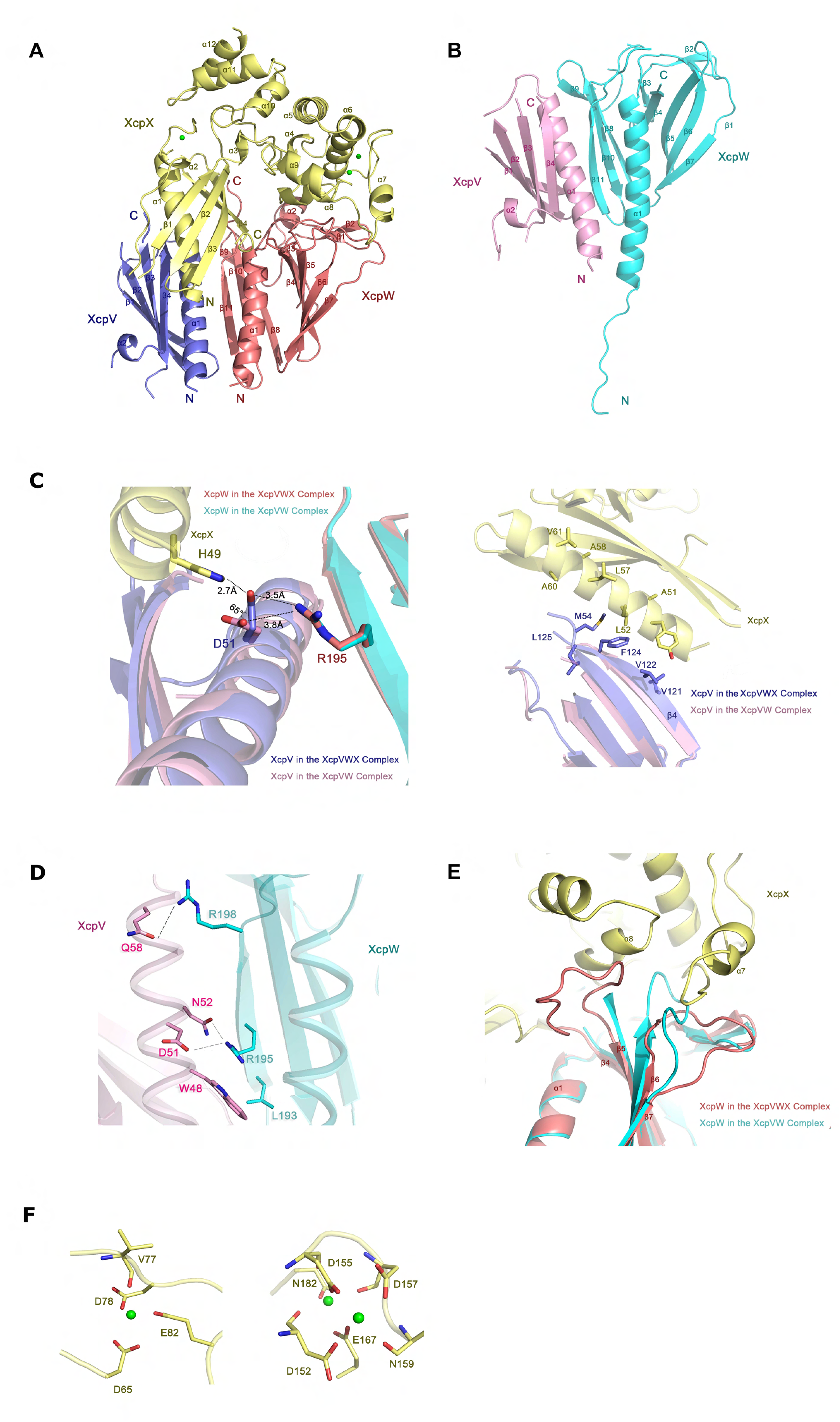
Crystal structures of XcpVWX and XcpVW complexes. (**A**) and (**B**) Overall structures of the XcpVWX and XcpVW complexes in ribbon representation. (**C**) The interaction interface shows that D51^(XcpV)^ is the critical residue to the formation of the XcpVWX ternary complex. (**D**) The interface between XcpV and -W reveals key residues involved in the interaction. (**E**) Comparison of the structures of XcpW in the complexes of XcpVWX (red) and XcpVW (cyan). The loop between β4 and β5 is stabilized in the XcpVWX complex by XcpX (yellow), and the loop between β6 and β7 is compressed by α7 of XcpX. (**F**) Three calcium-binding sites are found in XcpX. The newly identified calcium-binding site is shown on the left.

**Table 1.** Data collection and refinement statistics

|  | XcpVW | XcpVWX |
| --- | --- | --- |
| **Data collection** |  |  |
| Space group | $P2_1$ | $P2_12_12_1$ |
| Cell dimensions$\square\square$ | | |
| *a*, *b*, *c* (Å) | 40.11, 200.95, 66.45 | 61.5, 76.8, 102.9 |
| $\alpha$, $\beta$, $\gamma$ (°) | 90, 95.14, 90 | 90, 90, 90 |
| Resolution (Å) | 19.7-2.0 (2.1-2.0) * | 43.5-2.0 (2.15-2.0) * |
| $R_{\text{merge}}$ | 11.0 (76.3) | 4.7 (18.6) |
| $I / \sigma$ | 11.73 (2.77) | 10.3 (9.73) |
| Completeness (%) | 97.9 (100) | 99.2 (100) |
| Redundancy | 8.5 (8.7) | 9.8 (9.5) |
| **Refinement** |  |  |
| Resolution (Å) | 19.7-2.0 | 19.7-2.0 |
| No. reflections | 68806 | 31625 |
| $R_{\text{work}} / R_{\text{free}}$ | 20.62/25.80 | 19.09/24.81 |
| No. atoms | 8562 | 4300 |
| Protein | 8020 | 4015 |
| Ligand/ion | 20 | 3 |
| Water | 522 | 282 |
| *B*-factors |  |  |
| Protein | 36.3 | 36.3 |
| Ligand/ion | 35.1 | 35.1 |
| Water | 41.9 | 41.9 |
| R.m.s. deviations |  |  |
| Bond lengths (Å) | 0.008 | 0.008 |
| Bond angles (°) | 1.1 | 0.915 |

\* Values in parentheses are for the highest resolution shell.

* Values in parentheses are for the highest resolution shell.

The binding between XcpV (**Fig 1A, blue**) and -X (**Fig 1A, yellow**) is mainly established through the salt bridge between H49 ^(XcpX)^ and D51^(XcpV)^ (**Fig 1A, the left panel**). In the GspIJK structure, the similar interaction can also be found between W42^(GspK)^ and E45^(GspI)^ (**Figure EV2A**). Additionally, some nonpolar residues on the binding surface, *i.e.* L52, L57, and A60 on XcpX and M54, V122, and F124 on XcpV, introduce hydrophobic interactions to enhance the binding (**Fig 1C, the right panel**). The hydrophobic interactions draw XcpV closer to the N-terminus of XcpX in the ternary complex compared with the XcpV molecule in the XcpVW binary complex.

The main interacting contacts in this section of XcpVW include D51^(XcpV)^-R195^(XcpW)^, N52^(XcpV)^-R195^(XcpW)^, and Q58^(XcpV)^ - R198^(XcpW)^ as well as the hydrophobic interaction by W48^(XcpV)^-L193^(XcpW)^ (**Fig 1D**). The hydrogen bond between Q58^(XcpV)^ and R198^(XcpW)^ and the salt bridge between D51^(XcpV)^ and R195^(XcpW)^ substantially stabilizes the interaction between α1^(XcpV)^ and β11^(XcpW)^. The residue composition on the binding surface of XcpVW is not conserved compared with other species (**Figure EV2B, and 2C**). As the only residue in XcpV involved in binding both XcpW and -X (**Fig 1C, the left panel**), D51 experiences an angle change of ∼65° in the XcpVWX complex. Other contacts at the bottom of the bundled helices also maintain the binding of XcpV to -W (**Fig EV2D**). Interestingly, we observed a flexible N-terminal extension in XcpW, which acts as a linker between the transmembrane domain and 1^(XcpW)^ (**Fig 1B**). This unreported flexible structure is more likely to facilitate the assembly of the pseudopilus tip. Similar to the literature(Douzi et al., 2009), no significant interaction was identified between XcpX and -W. XcpX limits the spatial flexibility of the two loops on XcpW but forms no solid contacts (**Fig 1E**). One of the loop regions is between β4 and β5, and the other is between β6 and β7.

Our structure has revealed a novel calcium-binding site in XcpX, mainly constituted by D65, D78 and E82, in addition to the two canonical sites found in GspK of enterotoxigenic *Escherichia coli* (ETEC)(Korotkov & Hol, 2008) (**Fig 1F**). With further phylogenetic analysis, this binding site also exists in GspK^(XcpX)^ of some other Gram-negative species (**Fig EV3**). The calcium binding of XcpX was also confirmed by isothermal titration calorimetry (**Fig EV3A**). A systematic phylogenetic analysis of the full-length GspK demonstrated the evolutionary relationship between 37 different species of Gram-negative bacteria (**Fig EV3B, left panel**). Two main clades are clearly identified. The upper class branches into three subgroups, in which GspK of *P. aeruginosa* has closer relationship with *A. baumanii, A. borkumensis, P. alcaligenes*, and *P. stutzeri*; *L. pneumophila* is in the group with *B. pseudomallei, B. cepacia* and four *Pseudomonas* species. In the other group, GspK in *Vibrio* species are pretty close, and shares more similarities with the clade containing *S. boydii*, enterotoxigenic *E. coli*, enteropathogenic *E. coli, etc.* than the one that includes *Kelebsiella, Dickeya* and *Pectobacterium*. It is noted that GspK of some species in *Pseudomonas* genus, *e.g. P. putida*, which belong to the second group, are even closer to *E. coli* in evolutionary relationship. When specifically considering the sequences of the novel calcium-binding site, only the clade that includes *P. aeruginosa* has the third calcium binding site. No other clades share all the residues necessary for Ca^2+^ coordination (**Fig EV3B, right panel**), even those derived from the same main branch.

### Modeling the pseudopilus tip complex by SAXS

To characterize the entire structure of the pseudopilus tip complex and the spatial arrangement of the individual minor pseudopilins in the complex, we conducted SAXS to identify the structure of the tip complex of XcpUVWX in solution (**Fig 2, and Appendix Table S1**). The analysis of the scattering curve revealed that the molecular weight of this quaternary complex molecule is 79.5 kDa, in close agreement with the calculated complex molecular weight (80.3kDa), which rules out the possibility of aggregation or other oligomerization. The complex has a radius of gyration of 29.72. The curve of the paired distribution function, *p(r)*, suggested the maximum particle size (*D*_max_) of 95.04 (**Fig 2A**). Using DAMIN, the envelope of the entire the tip complex was reconstructed, and the crystal structure of the XcpVWX ternary complex and the modeled XcpU structure based on the homologous GspH of *E. coli* (PDB ID: 2KNQ) were docked into the envelope using CORAL (**Fig 2B**). The theoretical scattering profile was evaluated using CRYSOL, in excellent agreement with the experimental profile (◻^2^= 1.63).

**Figure 2.**
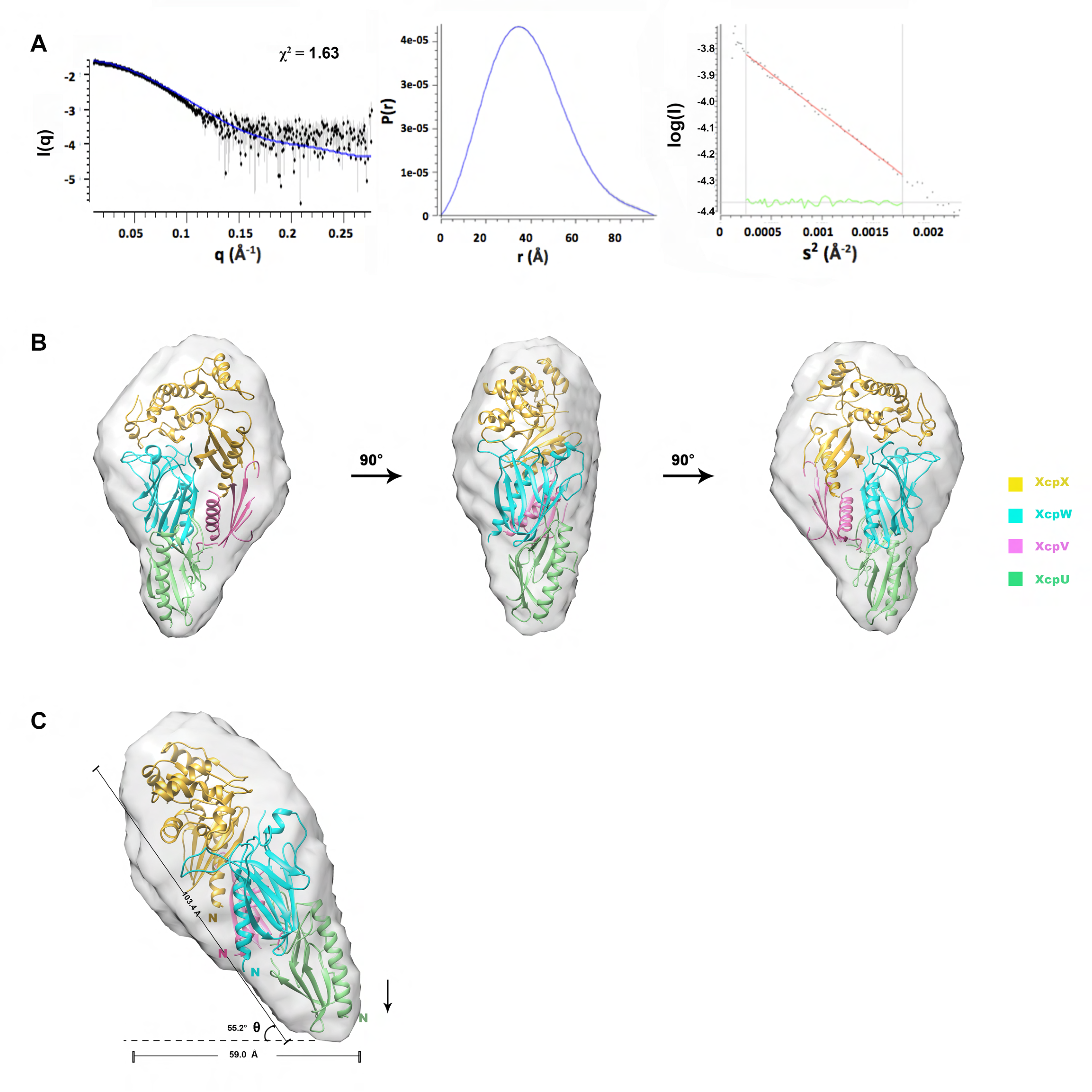
The architecture of the pseudopilus tip complex revealed by SAXS. (**A**) The SAXS data of the tip complex including the scattering data, the paired distance function and the Guinier plot. (**B**) The overall model of the tip complex in different angles. The crystal structures align with the SAXS envelope well showing that the binary complex of XcpVW links the molecules of XcpU and –X. (**C**) The spatial arrangement of the platform formed by the pseudopilus tip complex. The arrow directs the orientation of the N-termini of the four minor pseudopilins which all vertically point to the inner membrane.

The overall structure of the XcpUVWX tip complex adopts a ‘drumstick’ conformation according to the envelope (**Fig 2B**). In this model, XcpX is on the top (**Fig 2B, in yellow**) of the tip complex, and XcpU is at the bottom (**Fig 2B, in green**) with the critical XcpVW binary complex bridging in between (**Fig 2B, XcpV in pink, XcpW in cyan**). This overall structural arrangement is in line with the previously proposed assembly model of the tip complex(Douzi et al., 2009). Based on the previously proposed model of assembly(Cisneros et al., 2012a) that minor pseudopilins form the tip complex with their N-termini vertical to the inner membrane, our model shows that all the N-termini of the molecules are pointing down to the inner membrane (**Fig 2C**) with an angle (θ) of ∼55.2° between the horizontal plane and the slope of the tip complex. The span of the entire complex is 103.4 Å, and the length of the adjacent side of θ is ∼59 Å that is within the previously measured diameters of the outer membrane channels (55 Å – 64 Å) (Hay, Belousoff et al., 2017, Korotkov et al., 2011, Yan et al., 2017) to accommodate the tip complex.

### Compromising the core XcpVW binary complex abolishes Type II secretion

To identify the importance of individual minor pseudopilins in exoprotein secretion in *P. aeruginosa*, we compared the secretion pattern of *P. aeruginosa* PAO1 wild type (WT) with those of four minor pseudopilin transposon mutant strains with compromised pseudopilin expression (**Appendix Table S2**). Exoprotein secretion was abolished in the *xcpV* and *-W* mutant strains (**Fig 3A**), which is similar to the results in *K. oxytoca*(Cisneros et al., 2012a, Nivaskumar et al., 2016) and *xcpW*-deletion strain(Douzi et al., 2009). Compared with PAO1 and the *xcpU* and –*X* mutant strains, the secretion pattern of either *xcpV* or -*W* mutants showed the loss of two typical virulence factors secreted via the T2SS, namely elastase (LasB, ∼37 kDa) and lysyl endopeptidase (PrpL, ∼20 kDa), confirmed by mass spectrometry (data not shown). This indicates that the Type II secretory pathway was disabled in the *xcpV* and -*W* mutant strains, and that these two pseudopilins are essential to the T2SS.

**Figure 3.**
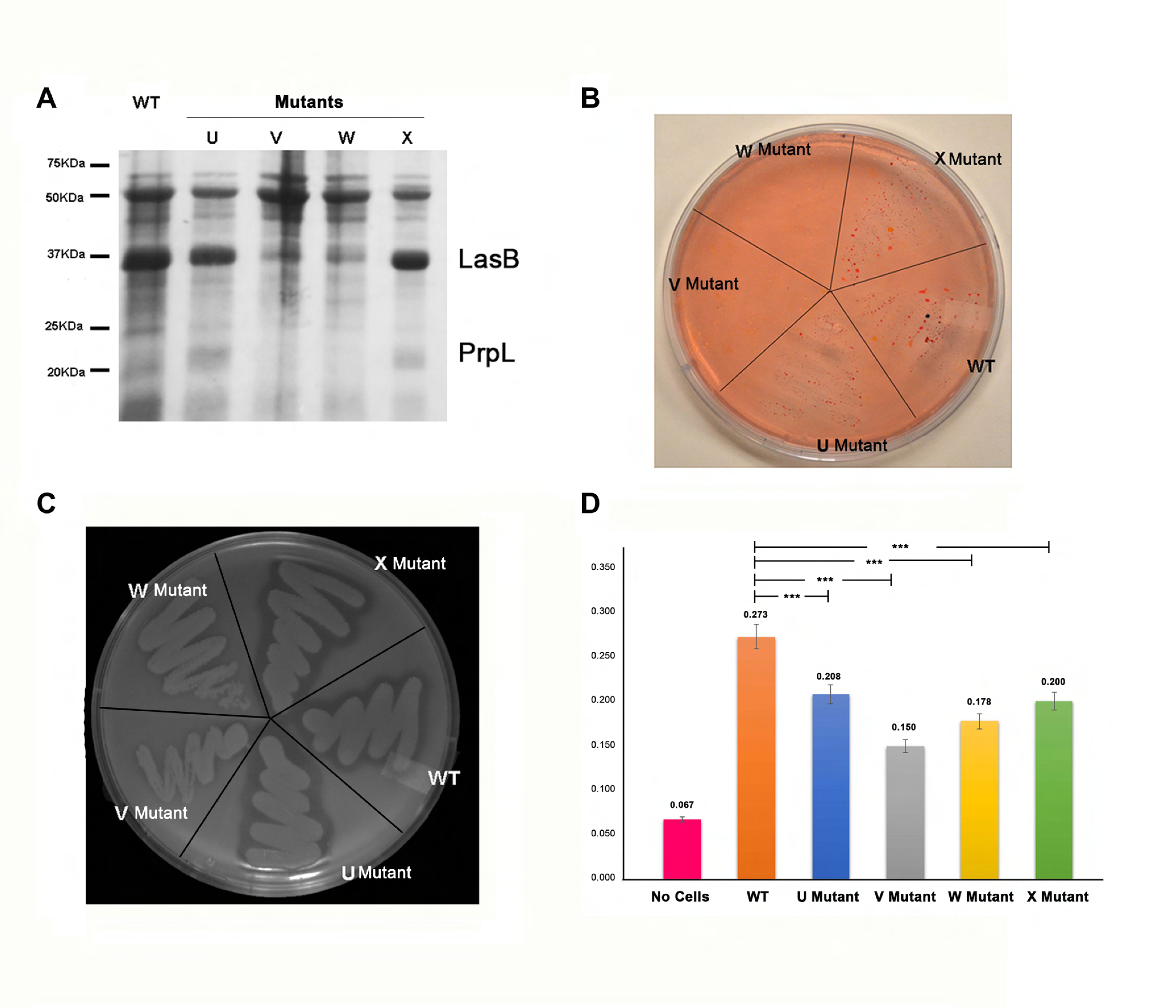
XcpV and -W form an indispensable core complex for the Type II secretion system. (**A**) Secretion analysis of WT or mutant strains. Compared with the WT, the secretion of the *xcpV* or -*W* mutant strains exhibited different expression patterns in which bands corresponding to secreted LasB (∼37 kDa) and PrpL (∼20 kDa) were significantly diminished. (**B**) Lipid agar assay of cell growth. The *xcpV* and -*W* mutant strains did not grow on the lipid agar plate, in contrast with the WT, *xcpU* and *xcpX* mutant strains. (**C**) Clearance of skim milk by secreted protease. The WT, *xcpU* and *xcpX* mutant strains generated clear rim by secreting protease while the *xcpV* and -*W* mutants did not because of the lack of the functional T2SS. (**D**) The activity of the secreted LasB indicates different levels of LasB secretion in different strains. *XcpV* and -*W* mutants showed less activities of LasB indicating a comprised T2SS.

Next, we assayed the strain survivability on lipid agar (**Fig 3B**). Only strains that secrete lipase to hydrolyze lipid, the sole carbon source, can survive under these conditions. The non-functional T2SS will therefore render cells nonviable. Neither the *xcpV* nor -*W* mutant strains survived on lipid agar, whilst the other two mutant strains (*xcpU* and -*X*) grew normally as the WT. Impaired expression of XcpV or -W also led to the disappearance of the secretion ring, created due to the proteolysis of skim milk by the secreted protease, on the skim milk agar (**Fig 3C**). Similar results were also observed when testing the activity of secreted LasB (**Fig 3D**). The supernatant of the *xcpV* mutant, containing secreted proteins, displayed the lowest LasB activity, suggesting that the secretion of LasB was significantly compromised. The supernatant of the *xcpW* mutant also had less LasB activity than the mutants of *xcpU* and -*X*.

Our secretion assays using transposon mutant strains are in accordance with the results previously reported that the loss of XcpV or -W impedes the function of the T2SS^21,22^. These findings reinforce the ideas that XcpV and -W form a core complex and act indispensible roles in the entire system. Though the Type II secretion is not equally disrupted in the *xcpU* and -*X* mutant strains as the *xcpV* and -*W* mutants, we still observed deminished LasB activities in *xcpU* and -*X* mutants compared with the WT (**Fig 3D**).

To validate the results of the *xcpV* and *xcpW* mutant strains, we performed a series of complementary assays by internalizing plasmids encoding the full-length DNA of *xcpV* and -*W* into corresponding mutant strains. Our results showed that the function of the T2SS was fully recovered (**Fig EV4**), suggesting that no polar effect occurred in our transposon mutants. The secretome, the survivability on lipid agar and skim milk clearance were rescued in the complemented strains compared with the original mutant strains. Taken together, these results fully support that XcpV and -W form a core complex indispensable to T2SS functions.

### Development of inhibitory peptides based on the structure of the interaction interface of the XcpVW complex

XcpV and -W form the core complex that bridges and incorporates XcpU and -X into the pseudopilus tip(Douzi et al., 2009, Korotkov & Hol, 2008, Yanez et al., 2008a). Therefore, we hypothesized that compromising this complex with inhibitory molecules would disrupt the tip complex and further inhibit the T2SS.

Results of MD simulation suggested that the upper region of the complex is the most dynamic for inhibitor penetration (**Fig EV5A and 5B**). Using this information and the interaction details illustrated by the complex structures, we respectively designed two mimicking peptides according to the sequences on either side of interaction interface. The peptides compete with the binding between XcpV and -W in this region, which protrude into the cavities on either side of the complex (**Fig 1D, 4A, and Fig EV5C**). While retaining all the interacting residues, we introduced several hydrophilic residues at noninteracting positions to enhance peptide solubility. The structure-based peptides were tested by MD simulation (**Fig 4B, and Fig EV5D**). Within 50ns of simulation, in the presence of the peptides, the initial interactions between XcpV and -W dissociated and new pairwise contacts formed during the simulation. Peptide 1 formed a small stable α-helix while Peptide 2 exhibited more flexibility. A rotation of 107.3º occurred to Peptide 1 after the simulation. Peptide 2 induced large conformational changes in the N-terminus of XcpV during the simulation.

**Figure 4.**
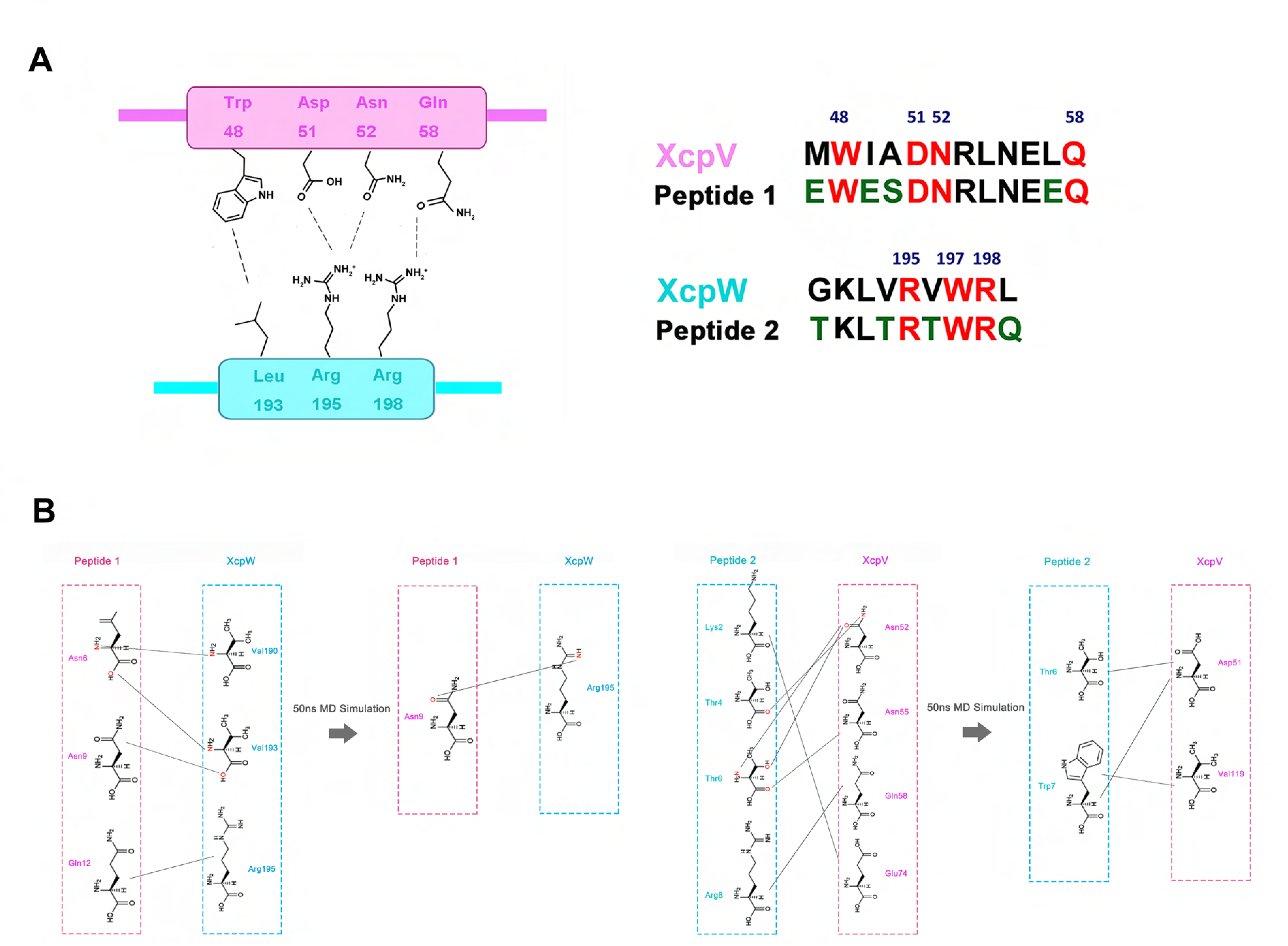
Inhibitory peptides are designed based on the interaction interface of XcpV and –W in the complex structure. (**A**) Based on the interface of the XcpVW binary complex (the left panel), two inhibitor peptides mimicking interacting segment of XcpV and -W respectively were developed (the right panel). All interacting residues (red) were retained while several hydrophilic residues (green) were introduced at non-interacting positions to enhance peptide solubility. (**B**) The re-establishment of the association of residues during the 50 ns of MD simulation of XcpV and -W in the presence of the structure-based peptides.

### Structure-based peptides disrupt pseudopilus tip complex and inhibit Type II secretion

To examine whether the structure-based peptides could compromise the formation of the XcpVW binary complex, we applied extrinsic fluorescence to detect conformational changes in the complex upon addition of the peptides. As the hydrophobic interactions among hydrophobic residues (*e.g.* W48^(XcpV)^-L193^(XcpW)^) account for the stability of the binary complex, it is expected that the more hydrophobic region in the interface is exposed to a fluorescent dye, the more fluorescence signals will be detected once the complex is compromised. Using Bis-ANS (4,4-dianilino-1,1-binaphthyl-5,5-disulfonic acid, dipotassium salt) as an probe, we added 5 μM and 10 μM of both peptides to 200 μM of the quaternary complex. The peptide-treated samples had an approximately twofold fluorescence increase of the samples treated with no peptide or the control peptide (NWKAKKHSLE) (**Fig 5A**), suggestive of an increased exposure of hydrophobic regions, probably due to the peptide-driven complex dissociation.

**Figure 5.**
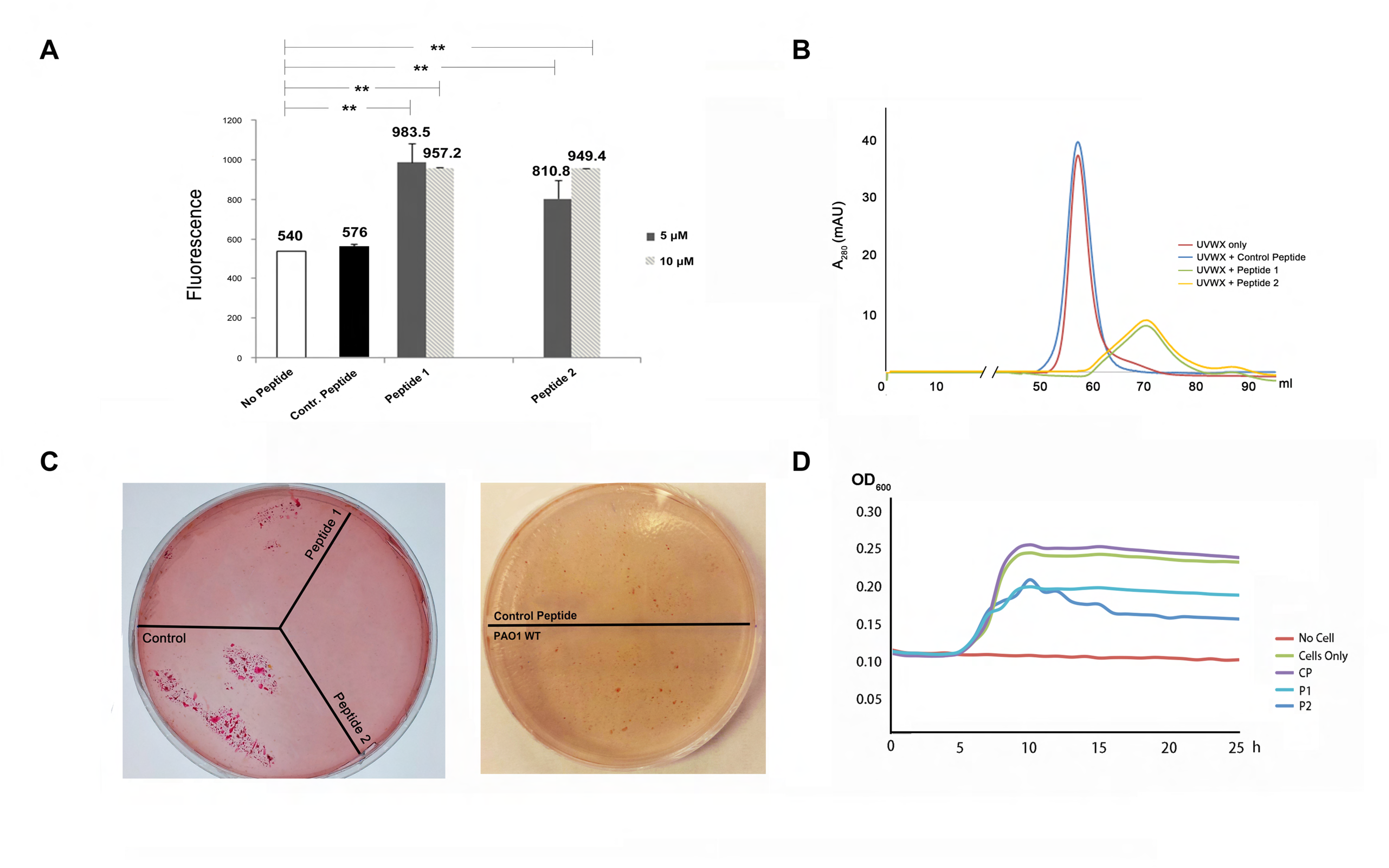
Structure-based peptides exhibited potency of inhibiting the Type II secretion system. (**A**) Fluorescence change of the XcpVW binary complex after incubation with peptides. Peptide treatment resulted in an increase in the fluorescence signal from Bis-ANS, suggesting the exposure of hydrophobic regions and complex dissociation. (**B**) The profiles of size exclusion chromatography showed different peak positions and heights in the absence and presence of the peptides. (**C**) Inhibition of lipase secretion by peptides. Bacterial survival is indicated by red colonies. Left panel: peptide-treated PAO1 cells failed to grow in contrast to untreated cells. Right panel: the scrambled peptide could not inhibit the lipase secretion, which showed no difference compared with the peptide-untreated cells. (**D**) Cell growth in lipid medium was hindered in the presence of the peptides. The PAO1 cells were treated by P1 and P2 while growing for 24 h in 96-well plate and the cell growth was inhibited compared with the cells that were treated with no peptide or the control peptide.

We further investigated the impact of the peptides on the integrity of the quaternary tip complex by monitoring changes in size exclusion chromatography (SEC). We incubated monomers of the four purified minor pseudopilins (**Appendix Fig S1A**) and subjected the mixture to SEC. A peak corresponding to the quaternary complex appeared, as confirmed by SDS-PAGE and analytical ultracentrifugation (**Fig 5B, red curve, Appendix Fig S1B and 1C**). Following the addition of the peptides, the quaternary complex disappeared in the chromatographic profiles and collapsed into a wider peak that displayed both decreased peak height (from ∼35 mAU to ∼10 mAU) and delayed peak position (from ∼58 ml to ∼70 ml), an evident sign of disruption of the quaternary complex (**Fig 5B, dark yellow and green curves**). However, no such disruption was found when the complex was treated with the control peptide (**Fig 5B, blue curve**).

*In vitro* evidence prompted us to assess the effects of the peptide on T2SS function *in vivo*. PAO1 cells, treated with or without the peptides, were plated on lipid agar, followed by the examination of cell viability (**Fig 5C, left panel**). Strikingly, compared with the WT strain without peptide treatment, little or no cells could grow on lipid agar after peptide treatment, demonstrating that the peptides effectively compromised the T2SS. By contrast, inhibition of lipase secretion did not occur when cells were treated with the control peptide (**Fig 5C, right panel**).

Furthermore, we verified the inhibitory effects of the two peptides on cell growth in the liquid lipid medium in 96-well plate. Using the OD600 of cells 24 h post inoculation for evaluating the extent of inhibition, the growth of peptide-treated cells from PAO1 was inhibited (**Fig 5D**).

### Structure-based peptides protect *C. elegans* from killing by *P. aeruginosa*

*C. elegans* has been widely used as the infection model for various pathogens(Huang, Li et al., 2014, Marsh & May, 2012, Powell & Ausubel, 2008). We thus attempted to use it to substantiate the inhibition of the T2SS in *P. aeruginosa* by the structure-based peptides. First, we examined how much the T2SS contributes to the virulence of *P. aeruginosa* towards *C. elegans*. *C. elegans* of L4 stage were infected with *P. aeruginosa* PAO1 WT and various transposon mutant strains. As shown in **Fig 6A**, we monitored the short-term (5 days) virulence of the *P. aeruginosa* strains towards *C. elegans*. The WT effectively killed *C. elegans* and over 70% of *C. elegans* infected with PAO1 died after 5 days post infection while *C. elegans* mixed with *Escherichia coli* OP50 (the control strain) remained alive. The transposon mutants of *xcpV* and -*W* as well as *lasB* largely attenuated the virulence of *P. aeruginosa* towards *C. elegans*, and only 15-25% of *C. elegans* died after five days. In comparison, *xcpU* and -*X* mutant strains resulted in only ∼20% less of *C. elegans* death rate than that caused by PAO1.

**Figure 6.**
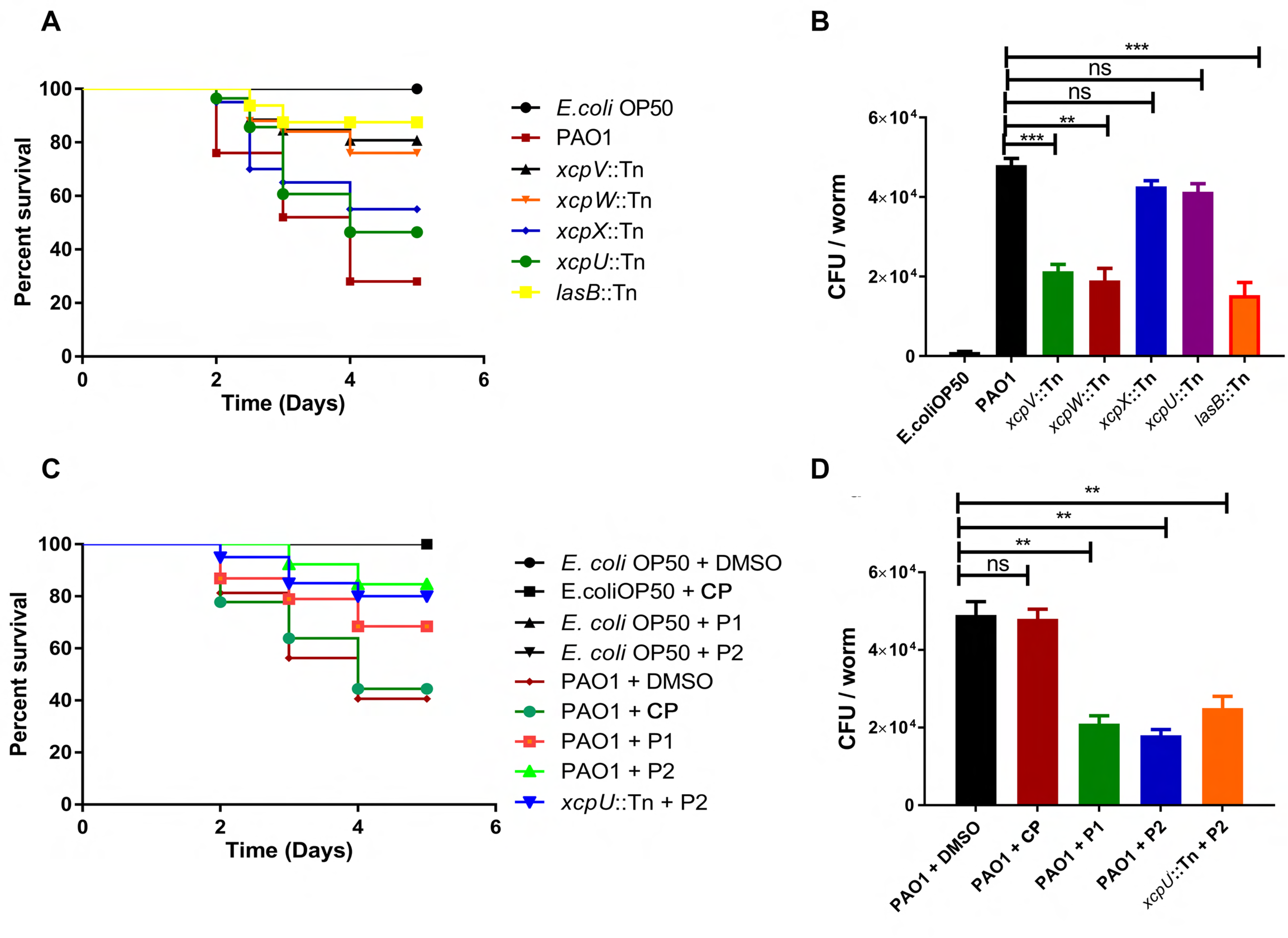
Effects of the structure-based peptides on *C. elegans* survival infected by *P. aeruginosa*. (**A**) The T2SS is critical to the virulence of *P. aeruginosa* towards *C. elegans*. The Kaplan-Meier plot for the survival of *C. elegans* with infection of PAO1 and various mutants indicates that the viability of *C. elegans* increased when infected by *xcpV*, *xcpW* and *lasB* mutant strains. (**B**) Bacterial CFU isolated from each infected worm. The reduction of CFU in *xcpV*, *xcpW* and *lasB* mutant strains indicates that the accumulation and virulence of *P. aeruginosa* have decreased. (**C**) Structure-based peptides increased the viability of *P. aeruginosa*- infected *C. elegans*. The Kaplan-Meier plot was established for *C. elegans* fed with either *E. coli* OP50 or *P. aeruginosa* in the absence or presence of structure-based peptides. ** P-value < 0.05 was adopted as statistically significant. ns: no significant difference. Structure-based peptides P1 and P2 have specifically affected virulence of *P. aeruginosa* and enhanced the viability of worms. (**D**) The level of accumulation of *P. aeruginosa* in *C. elegans* decreased in the presence of the inhibitory peptides.

The survivability of *C. elegans* may be closely related to *P. aeruginosa* accumulation in the guts of the worms.Therefore, we isolated and enumerated gut microbes. With similar growth rates in different strains (data not shown), the number of bacterial cells isolated from worms infected with the *xcpV*, -*W* and *lasB* mutants was 50% lower than the WT (**Fig 6B**). Consistently, the *xcpU* and -*X* mutants, though showing reduced cell numbers, exhibited no obvious difference from that of WT. These results reinforce the notion that the T2SS in *P. aeruginosa* contributes significantly to virulence towards *C. elegans*, and that XcpVW core complex is critical to the bacterial pathogenicity.

We continued to explore whether the blocking of the T2SS by our structure-based peptides could alleviate the virulence of *P. aeruginosa* towards *C. elegans*. Both peptides effectively protected *C. elegans* from being killed by the bacterium, as evidenced by the observation that *C. elegans* survival rate increased considerably from 40 % to 70% (P1) and to 85% (P2) within five days post infection (**Fig 6C**). Interestingly, P2 notably induced reduction of the virulence of the secretion-sufficient *xcpU* mutant, with the survival rate of *C. elegans* increasing from 40% (**Fig 6C, dark green line**) to 80% (**Fig 6C, blue line**).

The bacterial cell number in the gut of *C. elegans*, likewise, largely declined, when the pathogenic strains of PAO1 and the *xcpU* mutant were treated with the peptides (**Fig 6D**). More than twofold of cells (∼4.5 × 10^4^) were isolated from the worm infected with non-peptide-treated strains compared with the peptide-treated strains (< 2 × 10^4^), in which P2 (blue) showed more potency of inhibiting infection than P1 (green). Similar inhibition was also observed when the *xcpU* mutant strain was treated with P2. Apparently, the reduction of PAO1 cell number was not due to non-specific killing of bacteria by peptides, as they did not affect the number of *E. coli* OP50 in the intestine when *C. elegans* was mixed with OP50 (**Appendix Fig S2**).

## Discussion

The Type II secretion system is widely used in Gram-negative bacteria, including various pathogenic species, to secrete a vast range of virulence factors for infection. However, the molecular mechanism by which the T2SS secretes large, folded toxins has not been fully unveiled. Through the structural and functional studies, we have confirmed that the two minor pseudopilins, XcpV and -W, form a core binary complex that bridges the other minor pseudopilins to assemble the pseudopilus tip complex, which is essential for toxin secretion by the T2SS. Our findings in *P. aeruginosa* coincide with the results discovered in other species, *i.e. K. oxytoca*, that the pseudopilus tip is essential for the Type II secretory pathway. By showing that the lack of either XcpV or -W in the XcpVW complex resulted in the non-functional T2SS, we have provided experimental evidence that XcpVW binary complex plays an irreplaceable role in the T2SS of *P. aeruginosa*, which confirms the speculation that XcpVW functions as the core structure in the system(Douzi et al., 2009, Korotkov & Hol, 2008, Yanez et al., 2008a). Therefore, the destabilization of the core complex can lead to the inhibition of the T2SS.

Determination of the structures of the XcpVWX and XcpVW complexes of *P. aeruginosa* allowed us to define critical interacting residues/regions for developing inhibitory peptides. The structure-based peptides induced the destabilization and disruption of pseudopilin complexes *in vitro*, and obstructed the T2SS-dependent secretion of important virulence factors *in vivo*. Encouragingly, the inhibitory activities of these inhibitory molecules were confirmed in the infection assay of *C. elegans* by *P. aeruginosa*. The peptides, designed based on the structure of the core complex, specifically weaken the virulence of *P. aeruginosa* towards the worm. In conclusion, all our data have shown that the disruption of the core complex by the structure-based inhibitory peptide compromise T2SS function and reduces the virulence of *P. aeruginosa*.

Through SAXS, we have established the first structural model of the entire quaternary pseudopilus tip complex of the T2SS among all species, which provides important information to guide the study of the structure-related functions of the pseudopilus tip (**Figure 7**). From the overall model, the span of the length of the entire complex has been established. Based on the previously determined diameter of the channel bottom(Korotkov et al., 2011), to accommodate the tip complex, an angle between the bottom plane of the channel and the slope of the tip complex is required. By titling the tip complex relative to the inner membrane plane, not only the width of the overall structure would fit into the channel diameter but also all four N-termini would position perpendicularly to inner membrane, which provides further solid support for our SAXS structure. This model coincides with the previously proposed model of the assembly of the complex(Cisneros et al., 2012a) whereby the N-termini of the minor pseudopilins vertically insert into the inner membrane during assembly. Instead of grouping into a horizontal platform to push the secretion substrate, the tip complex forms a steep slope in which XcpU and -X lie on each sides of the XcpVW core complex. The recently-characterized structure of outer membrane channel of *P. aeruginosa* is more spacious than those of the other species, which provides sufficient room to allow the entering of the XcpUVWX tip complex(Hay et al., 2017).

**Figure 7.**
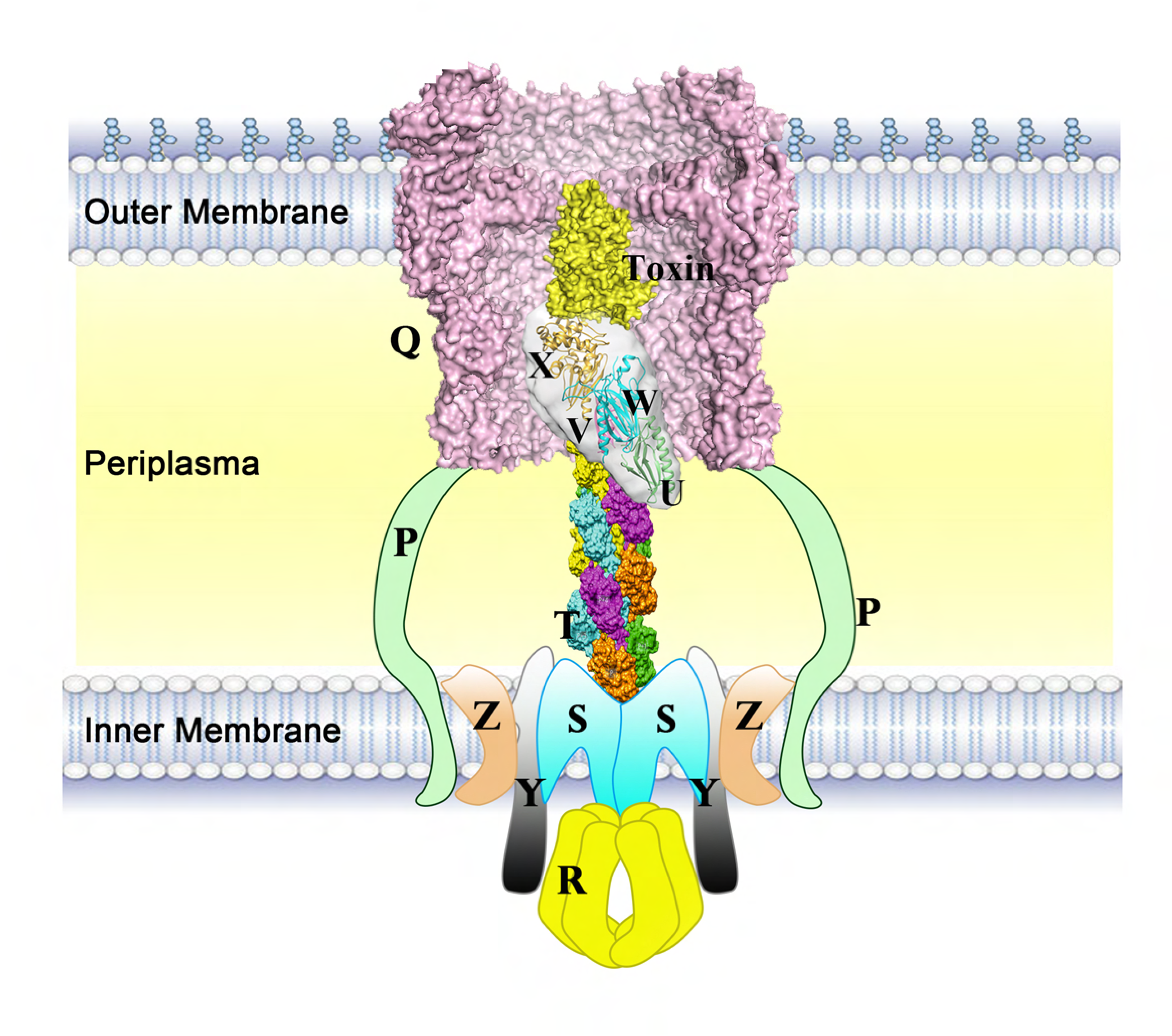
Schematic of the working model of the pseudopilus tip complex. The schematic shows the working state of the tip complex in the T2SS. The tip complex forms an angle with the horizontal plane during the polymerization of the major pseudopilin (adapted from the assembly model of PulG(Lopez-Castilla, Thomassin et al., 2017)) and enters into the outer membrane channel (PDB ID: 5WLN)(Hay et al., 2017).

The role of XcpU in the pseudopilus tip complex still needs to be elucidated, though its structure in *Vibrio cholera* has been elucidated(Raghunathan, Vago et al., 2014, Yanez et al., 2008b).Similar to the studies of *pulH* in *K. oxytoca*(Cisneros et al., 2012a, Possot, Vignon et al., 2000), the transposon mutant of *xcpU* showed almost no influence on toxin secretion.

Calcium is crucial for different secretion systems in bacteria. It is not only involved in the function of virulence factors but also necessary to the components of secretion systems(Brandhorst, Gauthier et al., 2005, Durand et al., 2005, Green & Mecsas, 2016, Kim, Ahn et al., 2005, Korotkov, Gray et al., 2009, Sarkisova, Patrauchan et al., 2005). We have identified a novel calcium-binding site in XcpX present in a small group of species (**Fig EV3B**). The calcium binding may be related to specific structural and/or functional properties of pseudopilus that awaits further exploration.

Protein structures have been applied to drug development(Anderson, 2003, Thiel, 2004, Zhang & Lai, 2011), in which the combination of combinatorial chemistry and structure-based design facilitate the development of diverse peptides after structure-orientated screening(Antel, 1999, Fosgerau & Hoffmann, 2015). Structure-based drug design has been used in different areas, including impeding virus infection, disrupting oncoprotein binding, inhibiting gene expression(Hernaez, Tarrago et al., 2010)’(Pazgier, Liu et al., 2009)’(Bartels, Schweda et al., 2007), *etc*. Our work, through both *in vitro* and *in vivo* assays, establishes the feasibility of using structure-based peptides to inhibit T2SS by specifically targeting the interaction interface of the XcpVW core complex, which significantly reduces the virulence of *P. aeruginosa*. As the first reported research on inhibition of T2SS by inhibitory molecules, our work may provide a new way for inhibiting bacterial secretion system and pathogenicity. This potentially presents an option for future therapeutic development to combat *P. aeruginosa* or even Gram-negative bacterial infections in general.

## Materials and methods

### Cloning, protein expression and purification

Soluble forms of individual minor pseudopilins (XcpU (29-172), XcpV (28-129), XcpW (28-203) and -X (29-313)) were constructed, using Kpn I and Xho I cut sites, into the pET32b vector which contains an N-terminal thioredoxin (Trx) followed by the TEV protease recognition sequence. Recombinant plasmids were introduced into BL21 (DE3) competent cells and protein expression was induced using 1 mM IPTG at 16 °C overnight after the OD600 of the culture reached 0.6. Harvested cells were sonicated for lysis using Buffer A (50 mM Tris, pH 8.0, 150 mM NaCl, and 10 mM imidazole), and cell lysate were obtained after high-speed centrifugation at 18,000 rpm for 30 min. The cell lysate was applied to Ni^2+^-NTA resins for incubation followed by washing the column with 50 ml of Buffer A. Proteins were eluted using Buffer A plus 200 mM imidazole. Eluted Trx-tagged proteins were subjected to overnight TEV protease digestion while dialyzing against Buffer B (50 mM Tris, pH 8.0, 150 mM NaCl, and 5 mM imidazole). Digested samples were reloaded to Ni^2+^-NTA resins to remove Trx tag and the flow-through that contains the pseudopilins was fractionated. Proteins were concentrated and loaded onto Superdex 75 column (GE Healthcare) for size exclusion chromatography using Buffer C (25 mM HEPES, pH 7.0, 150 mM NaCl).

To form the binary XcpVW complex, the purified components were mixed together in a molar ratio of 1.2:1 (XcpV: XcpW) and incubated overnight at 4 °C. The XcpVWX and XcpUVWX complexes were prepared in a similar way. The ratios between proteins were 1.5:1:1 (XcpV: XcpW: XcpX) in the XcpVWX complex and 1.2:1.5:1:1 (XcpU: XcpV: XcpW: XcpX) in the XcpUVWX complex. The complexes were purified by size exclusion chromatography. When XcpX was present, the buffer used for purification was Buffer D (25 mM HEPES, pH 7.0, 150 mM NaCl, and 1mM CaCl2). The formation of the quaternary complex was confirmed by analytical ultracentrifugation. One hundred μl of 100 μM of samples was added into the cell. The centrifugation was done at the speed of 1,000,000 g. The molecular size and properties of all fractions were analyzed.

For the T2SS recovery assay, full-length XcpV and -W DNA were inserted into the pUC18 vector with the cut sites of EcoR I and Xho I. The recombinant plasmids were internalized into XcpV and -W transposon mutant strains by electroporation following the standard electroporation protocol for *P. aeruginosa(Smith & Iglewski, 1989)*. Positive colonies were selected using 500 μg/ml of carbenicillin.

### Crystallization and structure determination

The XcpVW binary complex was crystallized at 20 °C in 10-15% PEG 3350, 0.1 M Tris, pH 7.0 and 0.5 mM CsCl using the method of hanging-drop vapor diffusion by adding 2 μl of 10 mg/ml of protein into 2 μl of condition solution. The XcpVWX ternary complex was first subjected to *in situ* digestion with chymotrypsin (protein: chymotrypsin = 100:1, w/w). Initial crystals were obtained from 19 %-22 % PEG 2000 MME, 0.1M Tris, pH 8.0-9.0, 0.2 M trimethylamine N-oxide by sitting-drop vapour diffusion at 20 °C and further optimized by microseeding. For cryo-protectant, 25% of ethylene glycol was used. Diffraction data were collected on beamline 23-ID-B at the Advanced Photon Source, Argonne National Laboratory (Argonne, IL). Complete data sets containing 360 frames of images were collected at 1.033-Å wavelength at 100 K from a single crystal with the exposure of 1 s per frame. The diffraction data were processed using XDS(Kabsch, 2010) and HKL2000(Otwinowski & Minor, 1997). The structure of XcpVW complex was solved by molecular replacement (Phaser(McCoy, Grosse-Kunstleve et al., 2007)) using EpsIJ (PDB ID:2RET) as a search model. The structure of XcpVWX complex was determined by Phaser using XcpVW (PDB ID: 5BW0) and GspK in GspIJK (PDB ID: 3CI0) as the search models. Structural refinement was carried out iteratively using PHENIX(Adams, Afonine et al., 2010) in combination with manual fitting using Coot(Emsley & Cowtan, 2004).

### Secretion Assays in *Pseudomonas aeruginosa*

All of the *Pseudomonas aeruginosa* strains were obtained from the *Pseudomonas aeruginosa* mutant library (http://www.gs.washington.edu/labs/manoil/libraryindex.htm)(Jacobs et al., 2003). The wild type *P. aeruginosa* PAO1 and pseudopilin transposon mutant strains were streaked onto LB agar and incubated at 37 °C overnight. Colonies of cells were picked and transferred to 10 ml of LB with additional incubation at 37 °C for 18 h. To harvest secreted exoproteins, 3 ml of cells were spun down at 15,000 rpm for 2 min, and the supernatant was mixed with 0.25 μl of saturated TCA to precipitate the proteins therein. Following two washes of protein precipitation with 250 μl of acetone, the isolated exoproteins were subject to SDS-PAGE characterization for visualization. The bands of proteins that were secreted in the WT, *xcpU* and -*X* mutant but not in *xcpV* or -*W* were cut and applied to mass spectrometry for further identification.

Lipid agar was prepared as described previously(Kagami, Ratliff et al., 1998). *P. aeruginosa* PAO1 WT or minor pseudopilin mutant strains were cultured at 37 °C overnight. Colonies were picked for further culturing in 5 ml of LB at 37 °C overnight and the resultant cultures were diluted by 100-fold when the OD600 reached 1.0 before streaking onto corresponding sections of lipid agar. Cells were streaked which were then incubated at 37 °C overnight followed by the observation of cell growth.

The skim milk plate was made following the published protocol(Durand et al., 2011). The *P. aeruginosa* PAO1 WT or minor pseudopilin mutant strains were prepared as stated above and cells were streaked on the skim milk agar followed by the incubation at 37°C overnight before observing and measuring the size of the secretion ring.

The elastase assay was conducted according to the description(Nakajima, Powers et al., 1979). An equal number of cells of different strains (∼500) was inoculated into the 96-well plate, and the supernatant of the overnight-cultured cells was obtained after spinning down cells at 13,000 rpm for 2 min. Pre-made 0.5mM of succinyl-Ala-Ala-Ala-p-nitroanilide (Sigma-Aldrich) was incubated with the supernatant for 5 min at 25°C and the reading at 410 nm was recorded.

For the secretion recovery assay, recombinant plasmids containing XcpV and -W were internalized into XcpV and -W mutant strains respectively by electroporation according to previous report(Smith & Iglewski, 1989). Positive colonies were selected using 500 μg/ml of carbenicillin, which were further used for secretome analysis, skim milk agar and lipid agar assays as described above.

### Isothermal titration calorimetry

XcpX calcium binding measurement using ITC was performed following the protocol as described(Radhakrishnan, Stein et al., 2009). The protein solution was loaded in the sample cell, and CaCl2 solution was loaded in the syringe. The concentration of XcpX ranged from 20 μM to 60 μM; it was titrated by CaCl2 with 10-fold in concentration at room temperature. The heat released per injection was measured, and all the data were analyzed using the Microcal Origin ITC software.

### Sequence searching, alignment and phylogenetic tree generation

Sequences of GspK proteins of different Gram-negative bacteria, including pathogenic and nonpathogenic species, were searched and obtained on Uniprot database (http://www.uniprot.org/). Using CLUSTAL Omega(Goujon, McWilliam et al., 2010, Sievers F), all the full sequences of GspK were aligned while maintaining gaps and output in the format of clustal with numbers. The scripts of tree file were also obtained which were further applied to drawing phylogenetic tree on iTOL (https://itol.embl.de/)(Letunic & Bork, 2016). An unrooted phylogenetic tree that reflects the relationship of GspKs in different species was generated by the iTOL server. The leaves on the same clade were classified into the same group, and different groups were separated in colors.

The aligned sequence was submitted to Espript 3.0 server(Robert & Gouet, 2014) (http://espript.ibcp.fr/ESPript/ESPript/) to analyze the similarities among different sequences. Referring to the results, the novel calcium binding site and the canonical calcium binding site were identified. The sequences of novel binding site were subjected to further alignment and phylogenetic tree generation.

### Small-angle X-ray scattering and construction of the structure of the tip complex

The XcpUVWX complex was prepared as described above, and concentrated to 6.15 mg/ml (calculated based on the absorbance at 280 nm) in a buffer containing 25 mM HEPES pH 7.0, 150 mM NaCl, 5% glycerol, 2mM DTT. SAXS data were collected at Cornell High Energy Synchrotron Source (CHESS, Ithaca, NY) combined with online size exclusion chromatography. Collected data and processed by RAW(Hopkins, Gillilan et al., 2017) and PRIMUS in ATSAS package(Franke, Petoukhov et al., 2017) to calculate the molecular weight, generate the Guinier plot, and paired-distance function. DAMMIF and DAMMIN(Franke & Svergun, 2009) was used for generating the *ab initio* bead model.

The model of XcpU was generated by Phyre2 server(Kelley, Mezulis et al., 2015) based on the structure of the NMR structure of GspH in *E. coli* (PDB ID: 2KNQ). The crystal structure of the XcpVWX complex and the modeled structure of XcpU were fit into the envelope by CORAL(Petoukhov, Franke et al., 2012), and further visualized in the interface of CHIMERA(Pettersen, Goddard et al., 2004). The reconstructed model was evaluated by CRYSOL in ATSAS to test the agreement with the SAXS profile.

### Peptide design and synthesis

Peptides were developed based on sequences of the upper region of the interaction interface from both sides of the complex. Peptide 1 mainly mimics the partial sequence of XcpV involved in the interaction (MWIADNRLNELQ). The important interacting residues were retained, including W48, D52, N52 and Q58. Similarly, Peptide 2 was designed based on the sequence of XcpW, in which R195, W197 and R198 were retained. To enhance the solubility, several hydrophobic residues were substituted by hydrophilic residues. Both peptides were synthesized by Biomatik. The peptide with a scrambled sequence of the interacting residues were also synthesized and used as a control.

### Molecular Dynamics Simulation

The interaction models of XcpV-XcpW, XcpV-P1, and XcpW-P2 were set up using the structure of the XcpVW complex as the starting conformation. Hydrogen atoms were added according to the standard protonation states at pH 7.0. The P1 and P2 models alone without proteins in the system were set up as a control in a similar way. All MD simulations were carried out using the AMBER9 package(D.A. Case, 2006) with a classical AMBER parm99SB (Cheatham, Cieplak et al., 1999, Jorgensen, Chandrasekhar et al., 1983) together with the parmbsc0 refinement(Perez, Marchan et al., 2007) and gaff(Wang, Wolf et al., 2004) force field parameters. The protocol for all MD simulations is described herein as follows: (1) systems were energetically minimized to remove unfavorable contacts. Four cycles of minimization were performed. Within each 5000-step minimization, harmonic restraints were applied on proteins, which decreased from 100 kcal•mol^−1^•Å^−2^ to 75 kcal•mol^−1^•Å^−2^, 50 kcal•mol^−1^•Å^−2^ and 25 kcal•mol^−1^•Å^−2^. The fifth cycle consisted of 5000 steps of unrestrained minimization before the heating process. The cutoff distance used for the nonbonded interactions was 10 Å. (2) Each energy-minimized structure was heated over 120 ps from 0 to 300 K (with a temperature coupling of 0.2 ps), while the positions of the proteins were restrained with a small value of 25 kcal•mol^−1^•Å^−2^. (3) The unrestrained equilibration of 200 ps with constant pressure and temperature was carried out for each system. The temperature and pressure were allowed to fluctuate around 300 K and 1 bar, respectively, with the corresponding coupling of 0.2 ps. For each simulation, an integration step of 2 fs was used. The SHAKE algorithm(Miyamoto & Kollman, 1992) was used on the bonds containing hydrogen atoms. (4) Production simulation runs of 50 ns were carried out by following the same protocol. A time point after thermal equilibration of 200 ps in each simulation was selected as a starting point for data collection. During the production runs, 15000-25000 structures for a simulation were saved for postprocessing by uniformly sampling the trajectory.

Energetics post-processing of single-trajectory/triplet-trajectory was performed for each MM-PBSA calculation by using the MM-PBSA module of AMBER9 program through molecular mechanics and a continuum solvent model. In MM-PBSA calculation, Gn_p/solve_ is nonpolar solvation free energy, which was calculated by using a solvent-accessible surface area (SASA) as follows:

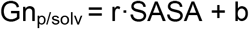

SASA is the solvent-accessible surface area, and was estimated using Sanner’s algorithm implemented in the Molsurf program in AMBER9 with a probe radius of 1.4 Å. The surface tension proportionality constant (r) and the free energy of nonpolar solvation for a point solute (b) were set to 0.00542 kcal mol^−1^•Å^−2^ and 0.92 kcal•mol^−1^, respectively.

For each of the three models, the last 20-ns trajectory of the production dynamics simulation was used for MM-PBSA binding free energy calculations, namely the 2000 snapshots of each model at a 10-ps interval for computation of enthalpy and 20 snapshots at 1000-ps intervals for computation of entropy. The following equation was employed to calculate the standard error (SE) of the binding free energy shifts:

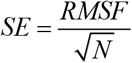

N is the number of snapshots collected in the calculations; RMSF is the root-mean-square fluctuation of the calculated ΔG binding values associated with all snapshots (Hao, Yang et al., 2010)

### Peptide disruption assays *in vitro*

For extrinsic protein fluorescence measurement, the XcpVW binary complex was purified as previously described and 5 μM or 10 μM (final concentration dissolved in water) of both peptides were added to 50 μM of complex samples (determined based on Beer-Lambert Law). The XcpVW binary complex was incubated overnight with and without the peptides. To each sample, 5 μM of Bis-ANS (Sigma) was added and incubated for 5 min. The fluorescence was detected by a Fluoromax-4 spectrofluorometer with the excitation at the wavelength of 385 nm, and the maximum emission signal was recorded at 515 nm.

The peptide disruption assay using size exclusion chromatography was carried out as follows. The peptides were added to the XcpUVWX quaternary complex in a molar ratio of 10:1 using Buffer D. After overnight incubation, samples with or without peptide treatment were loaded onto the Superdex 75 column for observing changes in the chromatographic profile.

### Peptide inhibition of Type II secretion of lipase

*P. aeruginosa* PAO1 WT cells were prepared for electroporation following the established protocol(Smith & Iglewski, 1989). Cells were diluted by 100-fold and incubated with each peptide at a final concentration of 100 μM. Electroporation was performed at 1600 V for 5 ms using the Multiporator (Eppendorf). Subsequently, 50 μl of aliquot of the cells was plated on the lipid agar plate for incubation at 37 °C for three days to observe cell growth on the plate.

For the assay in the lipid medium in 96-well plate, 10 μl of ∼1,000 PAO1 cells were added together with 2 μl of 10 mM of structure-based and control peptides into lipid medium with a total volume of 200 μl. The plate were incubated at 37 °C in the plate reader for 24 h with monitoring the cell growth. The OD_600_ values were recorded every 30 min.

### Cell culture and pre-treatment of *C. elegans*

*C. elegans* Bristol N2 WT was obtained from the Caenorhabditis Genetics Center (CGC, https://cbs.umn.edu/cgc/home). *Escherichia coli OP50* and *P. aeruginosa* were grown overnight at 37 °C in LB broth. *C. elegans* Bristol N2 was maintained at 23 °C on nematode growth medium agar plates seeded with *E. coli* OP50 using standard protocols(Smith, Laws et al., 2002). The peptides were dissovlved in water to make 25 mM and 10 mM stock respectively. The final working concentration of the peptides in the liquid medium was 250 μM.

### Nematode lethality assay

Bacteria from overnight culture were diluted at 1:100 and inoculated into 24-well microtiter dish containing 500 μL of liquid media (80% M9 and 20% BHI). The peptides were added when necessary at 1:100 dilutions from the stock solution. The L4 stage of *C.elegans* were washed with M9 minimal media and then pipetted (approximately 20 to 50 worms per well) into each well. Plates were incubated at 23 °C and scored for live worms every day. *E. coli* OP50 was used as a negative control.

### Detection of bacterial accumulation in nematodes

Bacterial accumulation assay was performed 24 hours after infection to determine the CFU count of PAO1 inside the exposed worms’ intestines. Infected worms were washed three times with M9 buffer and then maintained in the M9 buffer (1 mM sodium azide and 50μL/ml gentamycin) for ∼1 h to inhibit expulsion of bacteria from the worm intestine and kill any residual bacteria on the worms’ surface. Worms were washed three times with M9 buffer again. Approximately 10 worms were transferred to a 1.5 ml micro centrifuge tube and add 1 × PBS up to 100 μl. Four hundred mg of 1.0 mm silicon carbide particles were added to each tube and vortexed at maximum speed for 1 min to release bacteria. The lysate was briefly centrifuged and the supernatant containing bacteria was collected and plated onto LB agar plate for CFU count.

### Data availability

Coordinates and structure factors have been deposited in the Protein Data Bank under accession code 5BW0 for the XcpVW binary complex and 5VTM for the XcpVWX ternary complex.

## Acknowledgments

We thank the Canadian Institutes of Health Research, Natural Science and Engineering Research Council of Canada and Cystic Fibrosis Canada for their grant support. Dr. Bob Hancock kindly provided the mutant strains. Jaddie Ho is acknowledged for his technical help. Gregory Hicks has made great efforts to edit this manuscript. Z.J. is a Canada Research Chair in Structural Biology. W. Z and J. Z are supported by the Macau Science and Technology Development Fund (FDCT/066/2015/A2) and the Research Committee of University of Macau (MYRG2016-00199-FHS).

## Author Contributions

J.Y.Z., F.F and Z.J. devised experiments; J.Y.Z. crystallized XcpVWX complex and solved the structure, J.Y.Z. and N.N collected the SAXS data and constructed the model of the tip complex. F.F. crystallized and solved the structure of XcpVW complex. J.Y.Z. conducted all the secretion assays, and developed the peptide disruption experiments as well as the T2SS inhibition assay. W.Z. and J.Z. carried out *C. elegans* experiments. S.W. and M.W. performed the molecular dynamics and related calculation work. K.P. provided assistance in the *in vivo* assays. All authors contributed to the experimental design and assembly of the manuscript.

**Figure EV1.**
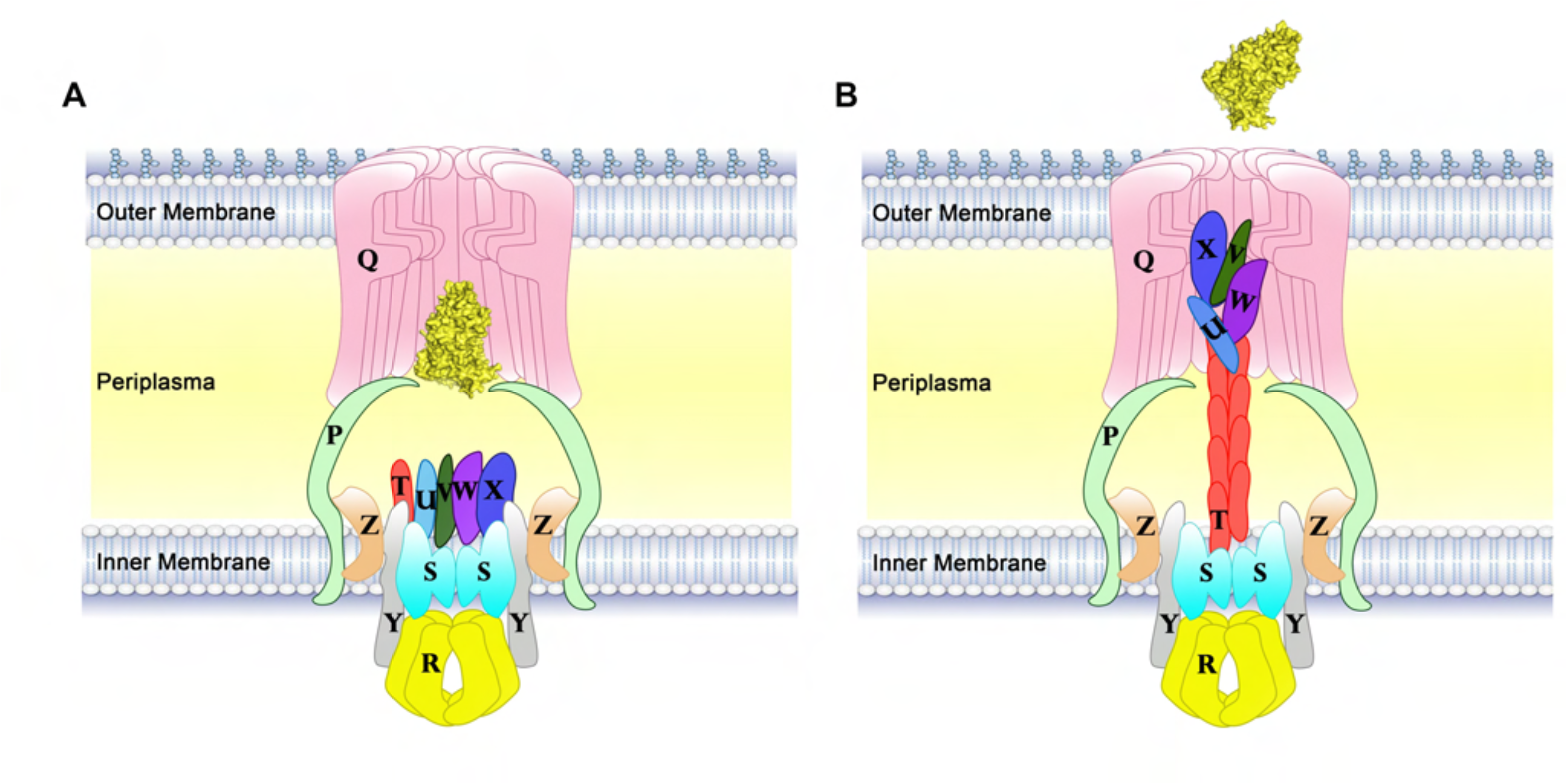
Schematic models of the Type II secretion system. (**A**) The Type II secretion system consists of multiple protein components that form several subcomplexes. (**B**) The working model of the secretion of virulence factors in T2SS. The minor pseudopilins form a quaternary pseudopilus tip complex, and the major pseudopilin XcpT polymerizes into piston-like pseudopilus body to secrete substrate (yellow).

**Figure EV2.**
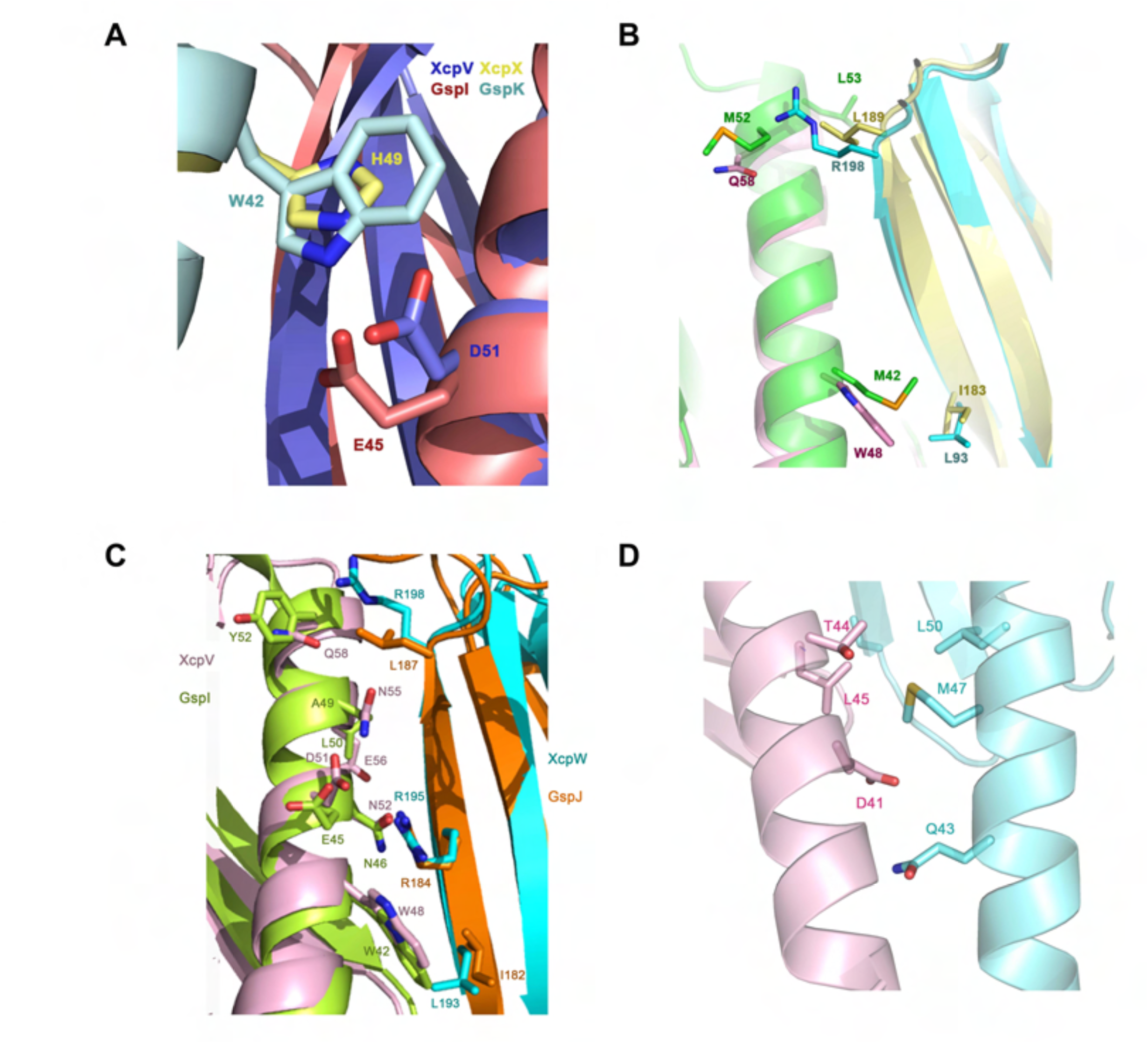
Protein-protein interactions in minor pseudopilin complexes. (**A**) Comparison of the formation of salt bridges in the XcpVWX (PDB ID:5VTM) and GspIJK complex (PDB ID: 3CI0). The salt bridge is the most important interaction between XcpV^(GspI)^ and XcpX^(GspK)^. In the XcpVWX complex (XcpV in blue and XcpX in yellow), the salt bridge is formed by D51^(XcpV)^ and H49^(XcpX)^ while in the GspIJK complex (GspI in red and GspK in cyan), E45^(GspI)^ and W42^(GspK)^ coordinate to build the salt bridge. (**B**) Comparison of important interacting residues between the XcpVW complex (XcpV in pink and XcpW in cyan) and the EpsIJ complex (PDB ID: 2RET, EpsI in green, and EpsJ in yellow). Q58^(XcpV)^ and W198^(XcpW)^ form a hydrogen bond to stabilize the interaction between XcpV and XcpW, whereas the hydrophobic interaction between M52^(EpsI)^ and L189^(EpsJ)^ contributes to the interaction. (**C**) Comparison of important interacting residues between the XcpVW complex (XcpV in pink and XcpW in cyan) and GspIJ in the GspIJK complex (GspI in light green, and XcpW in orange). (**D**) The residues involved in the interaction between XcpV and - W at the N-termini of the helices. In the XcpVW complex, XcpV and -W also interact through the contacts among residues in the bottom region of the main interface. Hydrogen bonding (D41^(XcpV)^-Q43^(XcpW)^) and hydrophobic interactions, *i.e.* L45^(XcpV)^-M47^(XcpW)^ and L45^(XcpV)^ - L50^(XcpW)^, are also involved in the binding.

**Figure EV3.**
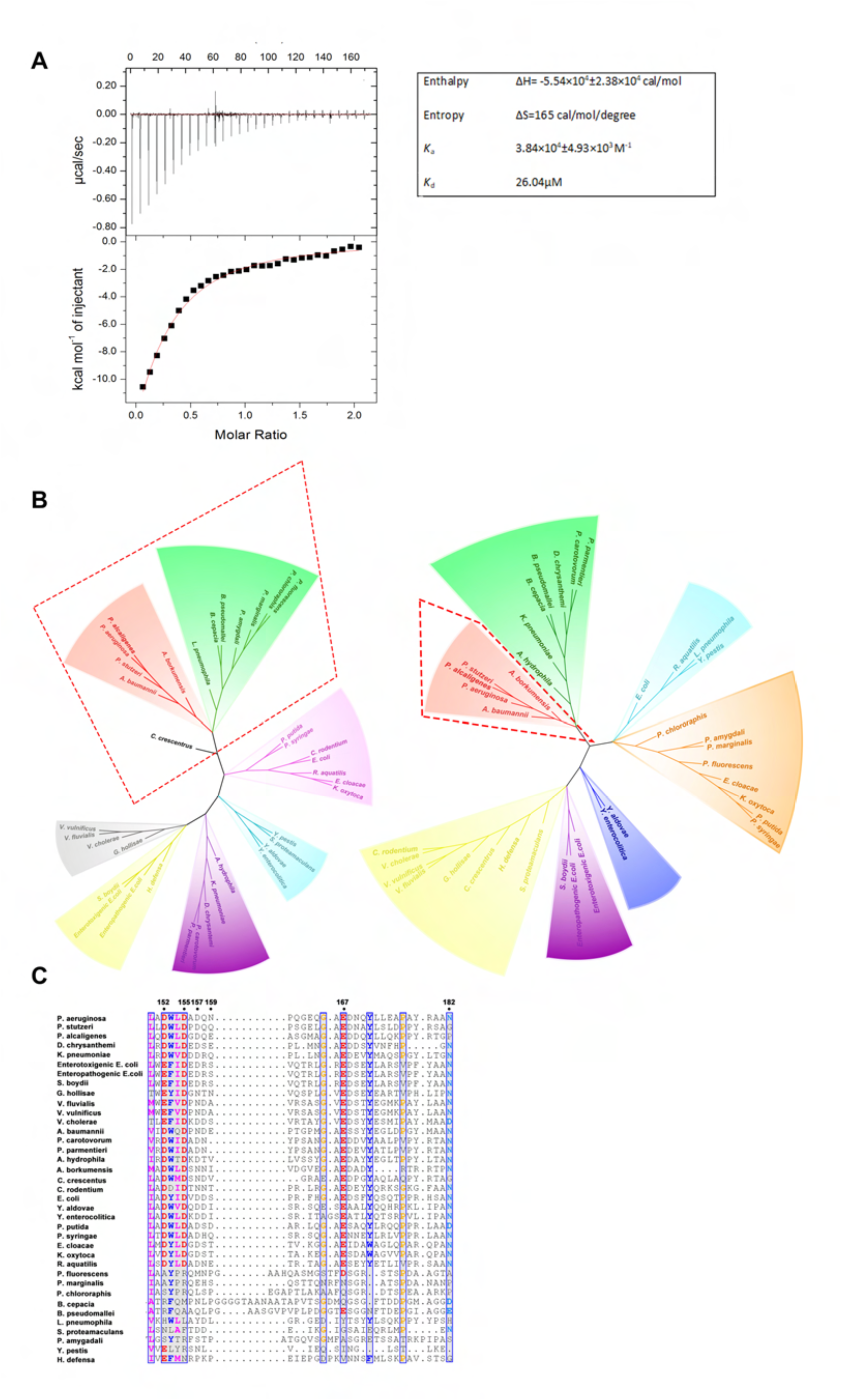
XcpX exhibits calcium-binding activity. (**A**) ITC results show that XcpX’s *K*_d_ of calcium binding is 26.04 μM. (**B**) Phylogenetic trees of GspK^(XcpX)^ proteins in different Gram-negative bacteria illustrate the evolutionary relationship. Left panel: phylogenetic trees of full-length GspKs with the first main clade boxed. Right panel: phylogenetic trees of the sequence of the new calcium-binding site in different GspKs shows that this additional calcium binding site only exists in limited Gram-negative bacteria (highlighted by the dotted box). (**C**) The alignment of the sequences of canonical calcium binding sites in different species implies that several species do not possess the ability of binding calcium. All the necessary residues for calcium binding are labeled.

**Figure EV4.**
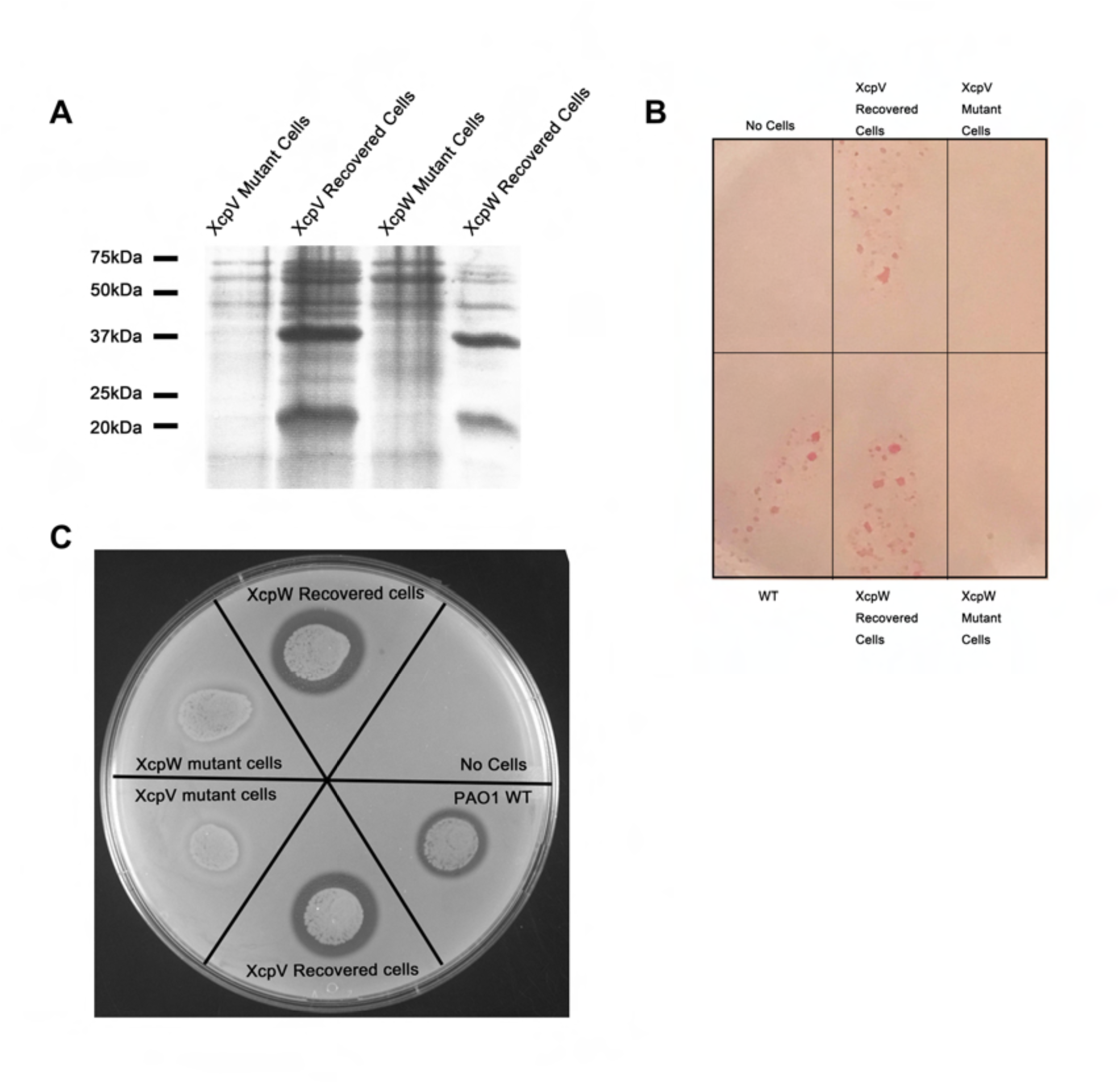
Complementary assays confirm that recovery of XcpV and -W expression in secretion-deficiency strains restored T2SS. Plasmids containing XcpV and -W were internalized back to xcpV or -W mutant strains respectively and restored the (**A**) secretion pattern of T2SS, (**B**) secretion of lipase, and (**C**) secretion ring of skim milk clearance, which indicate that T2SS has been recovered in mutant strains.

**Figure EV5.**
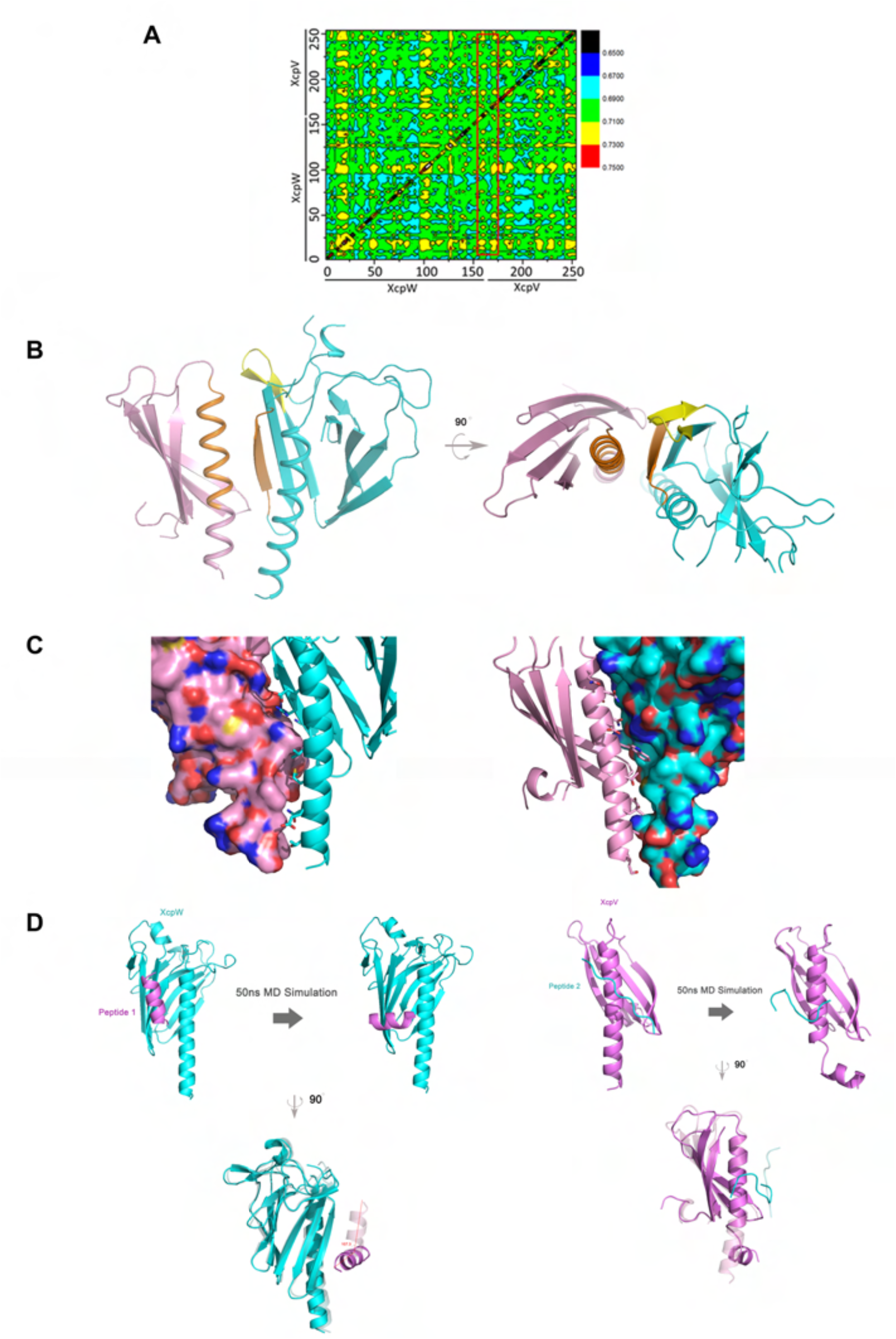
Molecular dynamics simulations of the XcpVW complex and the interaction between the peptides and pseudopilins. (**A**) Correlation factor reveals the most dynamic region in the XcpVW complex. The correlation factor of the XcpVW complex after a 50-ns MD simulation indicates the range of most dynamic residues (highlighted in the red box). During the simulation, all the key residues responsible for the interaction have experienced little change, maintaining stable binding. (**B**) Cartoon schematic shows that the dynamic region is located at the top of the complex (yellow), which suggests that the region could be targeted by inhibitory molecules. (**C**) Surface of the interaction interface in the complex. The interacting residues are shown in the interface for one protein component, while another protein is depicted in surface representation, and *vice versa*. (**D**) MD simulation (50 ns) of XcpV and -W in the presence of the peptides. Left panel: MD simulation (50 ns) shows the changes of the association between the peptides and pseudopilins. Binding of Peptide 1 to XcpW exhibits the binding free energy of −24.26 kcal/mol, which is weakened and further reduced to −0.92 kcal/mol due to angle change (107.3º). Right panel: during the simulation, the binding free energy changed from −34.26 kcal/mol to −24.13 kcal/mol. The flexibility allows Peptide 2 to influence the conformation of XcpV, which even induces conformational change at the N-terminus.

